# Diversity-generating retroelements in prokaryotic immunity

**DOI:** 10.1101/2022.06.02.494557

**Authors:** Ilya S Belalov, Arseniy A Sokolov, Andrey V Letarov

## Abstract

Adaptive immunity systems found in different organisms fall into two major types. Prokaryotes possess CRISPR-Cas systems, recognizing former invaders using the memorized (captured) pieces of their DNA as the pathogen signatures. Mammals possess a vast repertory of antibodies and T-cell receptors variants generated in advance. In this second type of adaptive immunity, a pathogen presentation to the immune system specifically activates the cells expressing matching antibodies or receptors. These cells proliferate to fight the infection and to form the immune memory. The principle of preemptive production of diverse defense proteins for future use can hypothetically take place in microbes too. In this study we test the hypothesis that prokaryotes employ diversity-generating retroelements to prepare defense proteins against yet-unknown invaders. We identified several candidate defense systems and characterised them.

## Introduction

The only known type of adaptive immunity in prokaryotes is CRISPR-Cas systems [1]. The basic principle of the CRISPR-Cas systems is memorizing invaders using fragments of their DNA, captured as so-called spacers into CRISPR arrays. The RNA transcripts of these arrays in the form of guide RNA target the interfering proteins to the DNA or RNA of invaders carrying the sequences of the spacers. Vertebrate immune systems, on the other hand, generate a gigantic pool of immunity sensors (antibodies and T-cells receptors) before contact with any pathogen [2]. Pathogen invasion induces the proliferation of the cells capable of producing antibodies or receptors that fit the pathogen antigens. These activated immune cells fight the pathogen and form the immune memory, facilitating a more rapid response if the pathogen returns. Among the means of fine tuning the immune sensors in the activated lymphocyte clones, vertebrates have developed the mechanism of somatic hypermutation. Hypermutation, mediated by specific mechanisms, is known in prokaryotes as well [3, 4]. However, there is no known defense system employing an analog of somatic hypermutation in bacteria and archaea. We hypothesize adaptive prokaryotic immunity systems, which employ diversity-generating retroelements (DGRs) in order to produce diverse antiviral proteins.

DGR systems include three essential components: reverse transcriptase (RT), template repeat (TR), and variable repeat (VR) [5]. The sequences of TR and VR are very similar to each other. Hypermutation occurs during RT-mediated cDNA synthesis from transcribed TR. Mutated TR copy replaces homologous VR in a process referred to as retrohoming. Such events occur independently in different cells creating a large diversity of the VR variants at the population level. The utility of the VR alterations depends on the function of the VR-containing protein [5–7]. In the *Bordetella* phage BPP-1, the archetypal DGR system [8], the VR is located within the major tropism determinant (*mtd*) gene, encoding the tail fiber protein. Multiple mutations in this protein enable adaptation to the host phase variations of the surface molecules *via* the viral tropism switch. Our hypothesis suggests that it may be possible to find hypervariable genes in prokaryotes constructed from VR with functions in antiviral immunity. This defense system would then play a role in producing preadaptation to new (or mutated) viruses.

The defense machinery employed by microorganisms includes a vast variety of proteins and protein systems [9, 10], the majority of which are likely yet to be described. Different types of defense systems are often co-localised in genomes, forming so-called ‘immunity islands’ [11]. This association has created a basis for successful identification of new systems of prokaryotic defense [12, 13]. Using an analogous approach, in this study, we test the hypothesis that prokaryotes organize defense mechanisms around DGR systems.

## Results

We aimed for comprehensive identification of new prokaryotic defense systems, utilizing DGRs, in the microbial pangenome, including bacteria and archaea. For each of the 32 321 publicly available DGR-encoding nucleotide sequences [6, 14, 15], we explored the vicinity of each identified DGR component (TR, VR, and RT) for the genes encoding immunity-related protein families (Pfams). For each Pfam occurring within 10 kbp from the abovementioned DGR components (1 348 709 proteins and 4907 Pfams in total), we computed the Icity score [16], a metric reflecting the probability of functional association between the given Pfam and the components of DGR systems (see Methods). High Icity score values occur only for a fraction of Pfams (see histogram in Fig. 1), 617 of which had Icity scores > 0.7. These Pfams fell into three types: (1) Pfams containing the VR sequence(s), (2) domains fused to RT, and (3) accessory proteins potentially functionally coupled to the DGR system. We hypothesize that any of these types may be involved in a DGR-dependent prokaryotic defense machinery.

**Fig. 1.**
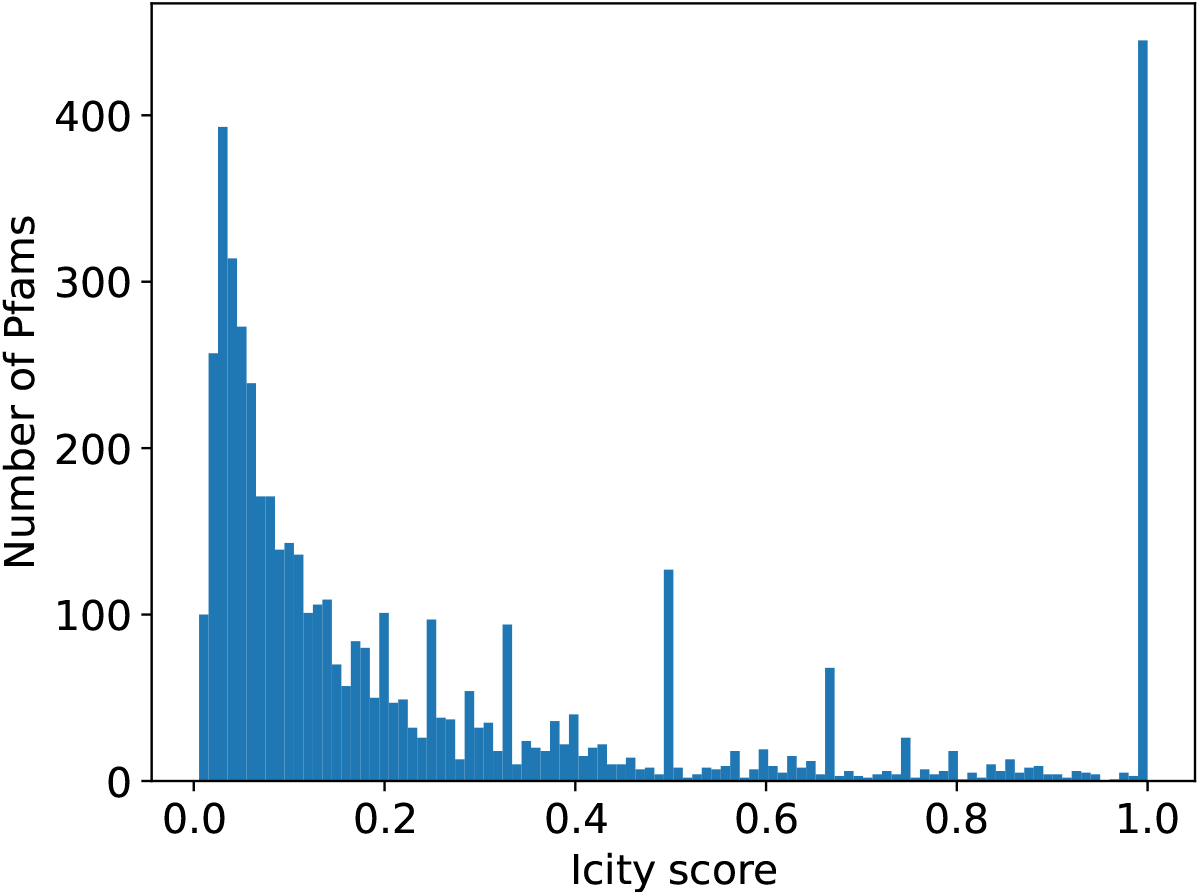
Icity score distribution among Pfams found within 10 kbp from components of DGR systems.

Prokaryotic immunity systems often co-localise, forming so-called ‘immunity islands’ [11–13]. A catalog of the Pfams found in genomic neighborhoods of known immune systems and therefore potentially involved in prokaryotic antiviral defenses was recently built [12, 13]. Further, in this paper, we will refer to such Pfams as ‘immune Pfams’. We further focused on DGR systems functionally coupled to immune Pfams. In total 8 557 out of 32 321 (27%) analyzed DGR systems had in their vicinity an immune Pfam with a high Icity score. For further analysis, we kept only sequences possessing at least one immune Pfam and presumably functional DGR components. We classified all sequences according to Pfams similarity. Fig. 2 shows Pfam compositions which occurred more than 26 times in our dataset. In Fig. 2 one can see that several compositions are subsets of the other ones (with larger Pfam numbers). The larger compositions (supersets) can occur both more frequently and less frequently compared to the occurrence of the subsets found separately. These composition variants may represent different (sub)types of DGR-based immune systems, or our observation may reflect the fact that about 50% of the selected sequences are shorter than 20 kbp (see Fig. 3). The maximum distribution at 3 kbp. So Pfam compositions appearing as subsets may be encoded by shorter sequences, lacking the rest of the Pfams from the corresponding superset.

**Fig. 2.**
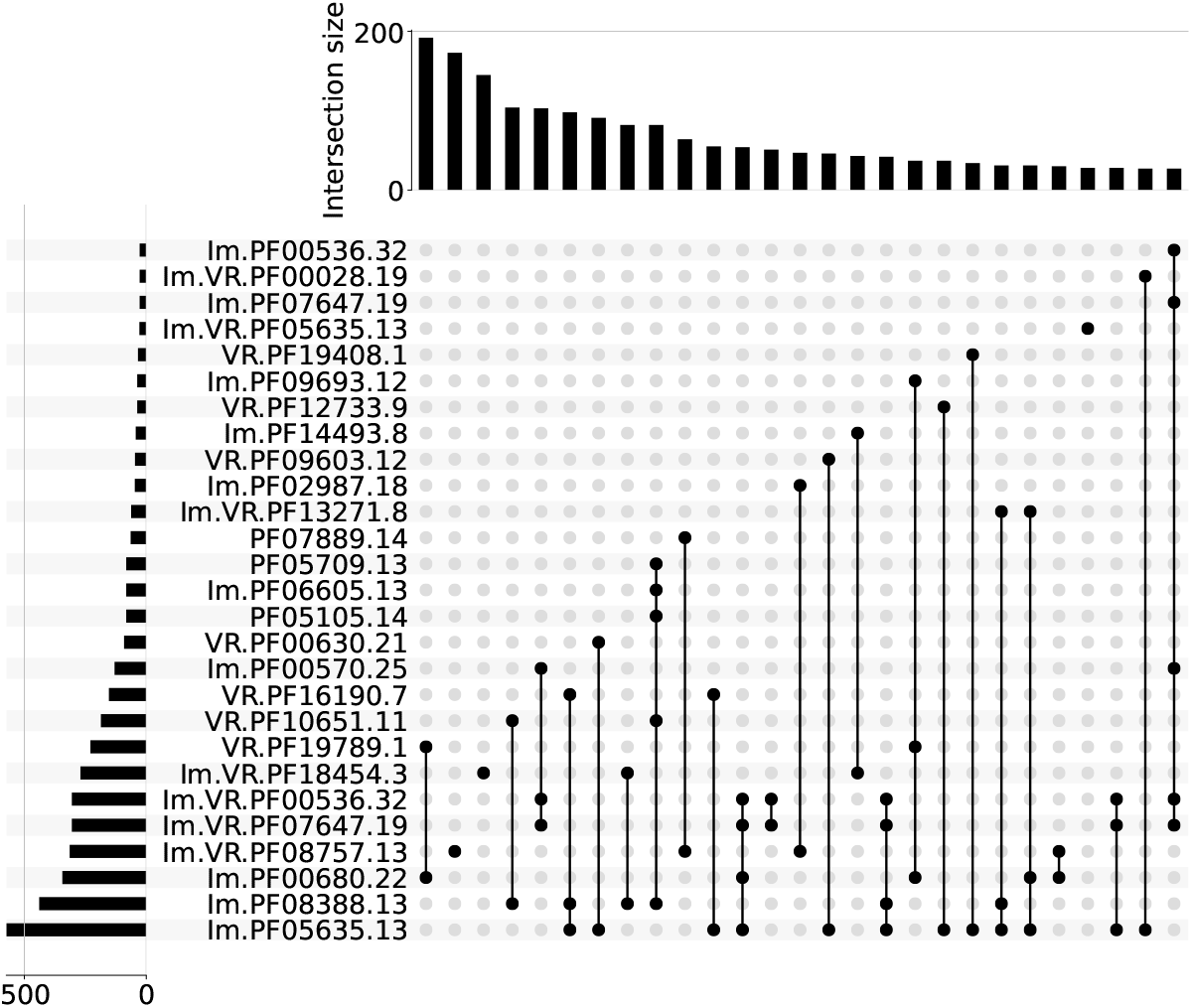
Pfam compositions with high Icity score and including at least one immune Pfam. Pfams associated with known prokaryotic defense systems have the prefix ‘Im.’. Proteins containing Variable Repeats have the prefix ‘VR.’.

**Fig. 3.**
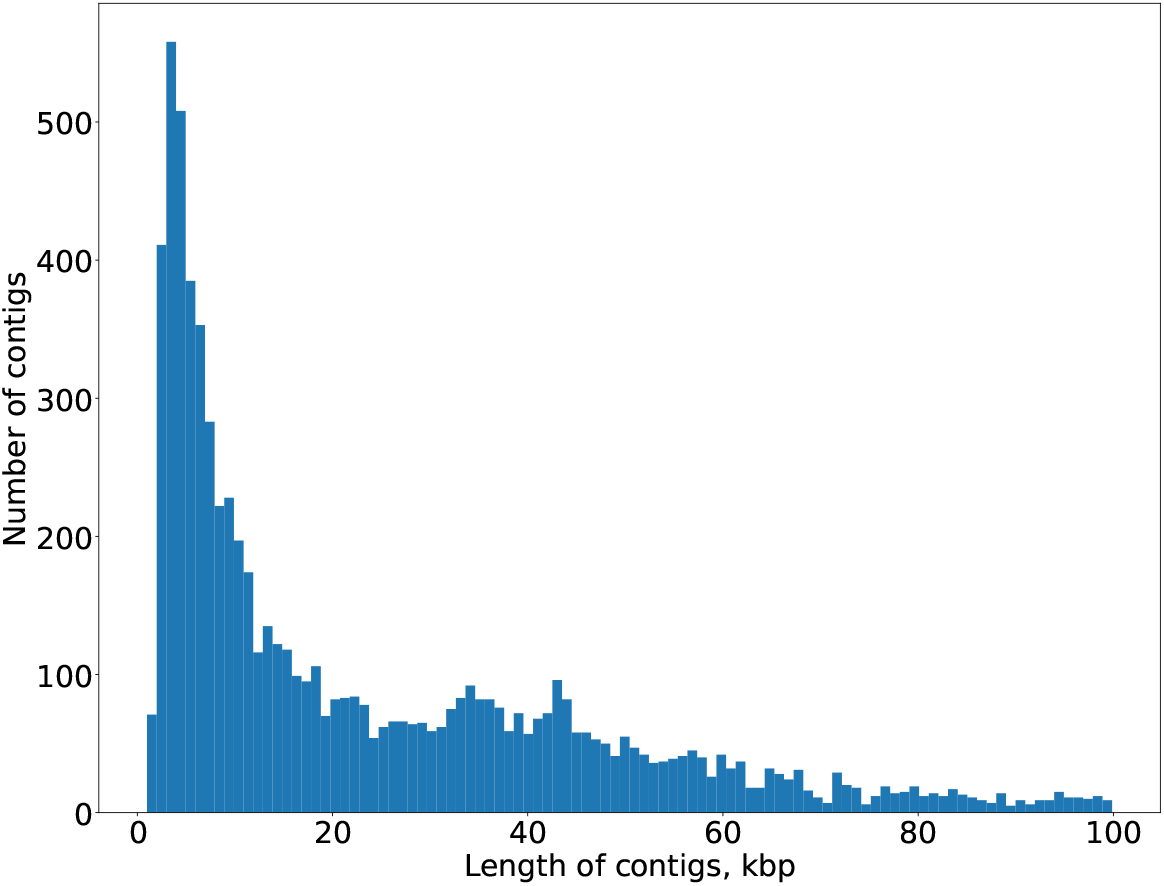
Contig lengths distribution.

Using complete genome sequences would be preferable in order to maximize the chance of observing intact and untruncated DGR-based immune systems. However, the availability of only a small number of such sequences (complete genomes encoding DGR) undermined our effort to find potential immune systems utilizing DGR mutagenesis. Thus, including longer contigs in our analysis presented a reasonable trade-off in this context. When we visualized Pfam compositions encoded by contigs longer than 20 kbp, we obtained a qualitatively similar picture, *modulo* halved dataset (Fig. 4). Supersets of both types (having more common or rare subsets) were preserved, occurring in a relatively uniform manner across the distribution of Pfam compositions. For each subsequent analysis presented here we considered only compositions encoded in more than 25 sequences, among which at least four contigs had a length > 20 kbp. Eighteen compositions in total satisfied these criteria.

**Fig. 4.**
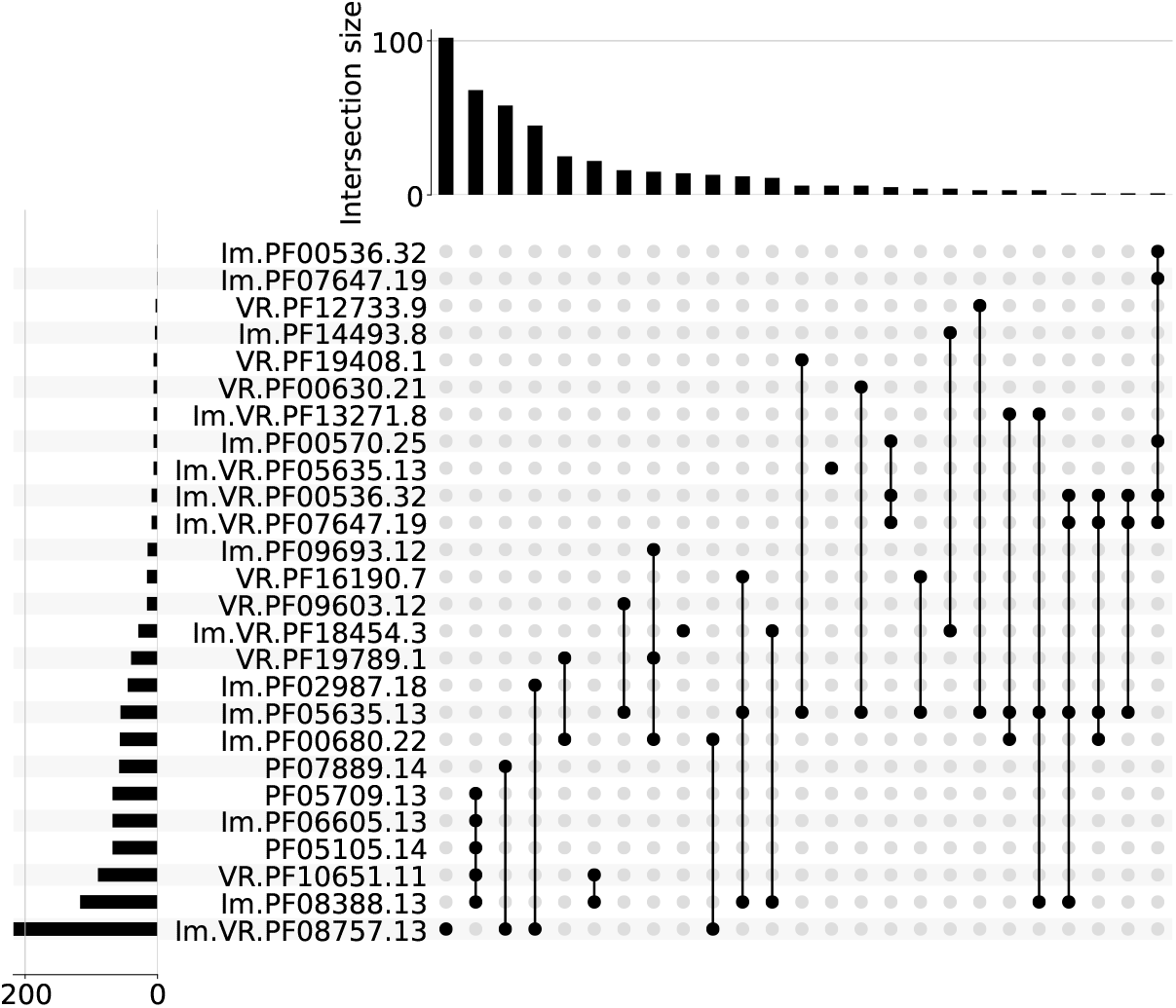
Pfam compositions from Fig. 2 among contigs longer than 20 kbp.

Prokaryotic defense systems are often spread via horizontal gene transfer (HGT) [11]. Nevertheless, the events of horizontal gene transfer are relatively rare. Host adaptation to a laterally acquired genetic module (*e*.*g*. a defense system) should include, among other factors, gradual bringing of the codon usage frequencies of the acquired module closer to the new host. Thus, the disparity in codon usage bias between a putative gene system and the rest of the genome may indicate a recent HGT event. Fig. 5 shows the proportion of sequences, for which the Kolmogorov – Smirnov test resulted in significant p-value (< 0.05), for eighteen selected Pfam compositions.

**Fig. 5.**
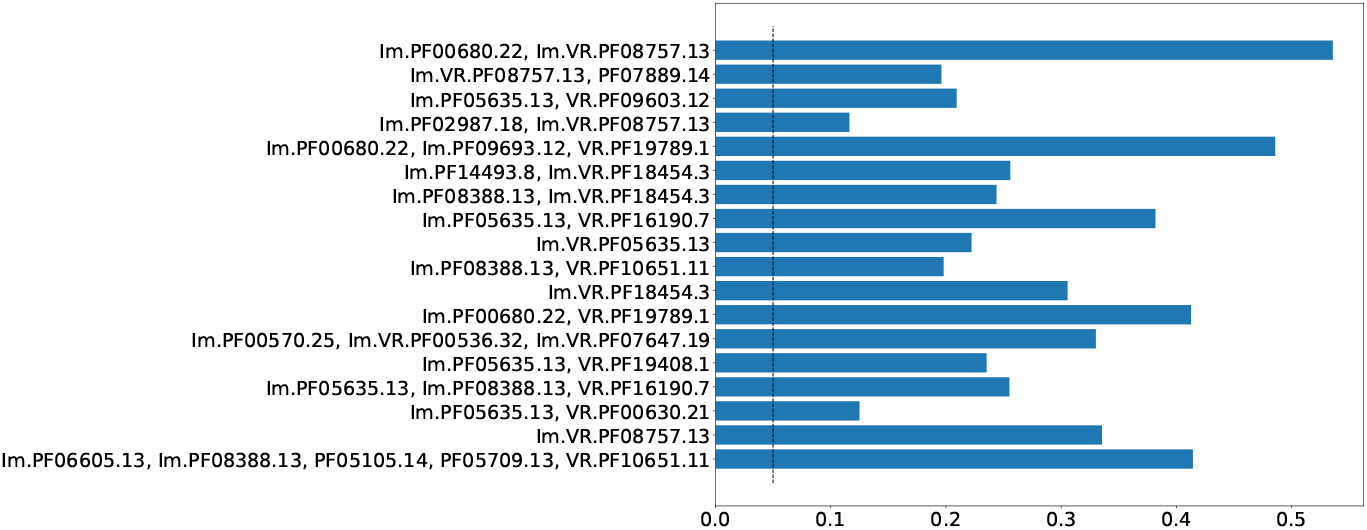
Proportion of sequences, for which codon usage bias in the genes in close neighbourhood (*±*10 kbp) was significantly different (KS p-value < 0.05) from most remote part of the same size of the same sequence. Dashed line indicates value of 0.05.

Prokaryotic defense systems also get adopted and repurposed by viruses, *e*.*g*. CRISPR-Cas systems which have been found in bacteriophage genomes [17] and R/M systems [18]. However, phages typically possess defense or antidefense mechanisms distinct from the systems found in bacterial genomes [19]. Although HGT between prokaryotes and their viruses has been documented many times, the functionality and organization of a defense system rather make it suitable either for cells or for viruses. Thus, for each type of putative prokaryotic immune system one should expect uneven distribution between cellular and viral genomes. Fig. 6 shows the proportion of cellular *versus* viral (including prophages) sequences encoding putative DGR-based defense systems.

**Fig. 6.**
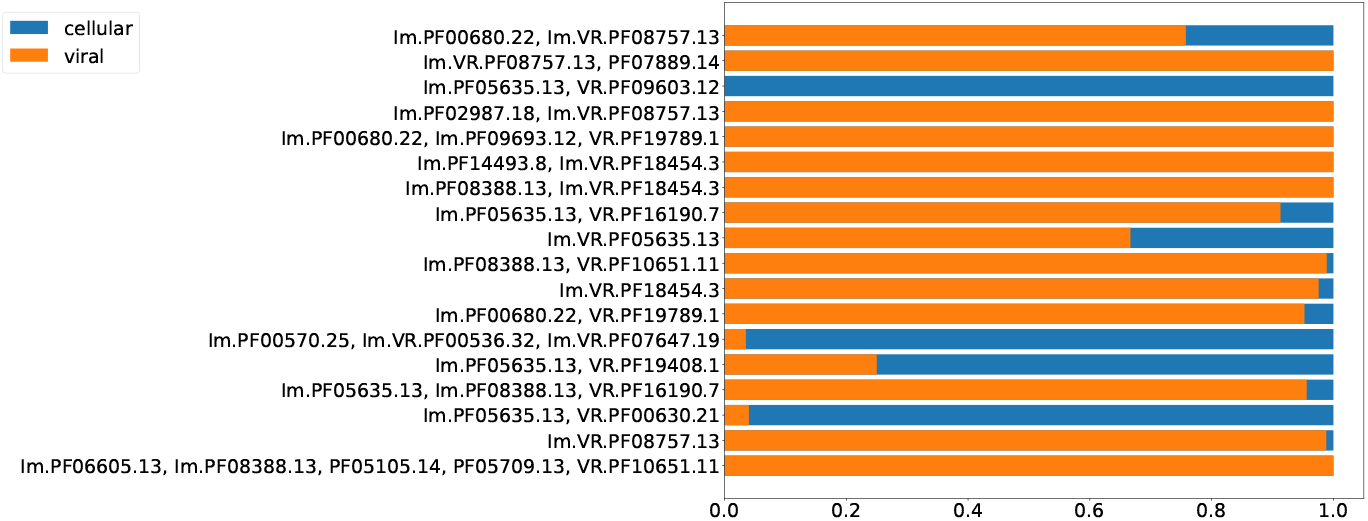
Proportion of viral and cellular residence for selected Pfam compositions.

The eighteen selected putative DGR-based immune systems share only one predicted protein functional category – reverse transcriptase activity. The reverse transcriptases of DGR systems are classified into six major phylogenetic clades [6]. By analogy with the data on CRISPR-Cas systems [20] and other prokaryotic defense machinery [21] one would expect that most of the aforementioned types of putative DGR-based immune systems have monophyletic origin. Therefore, each type should possess a single type of the RT. Presence of a second type of reverse transcriptase within a single type of DGR system is in theory possible (due to the lateral transfer events) but is less likely in our hypothesis. And, in the case of polyphyletic origin of a particular type of DGR-based system, accessory proteins would be better suited for one type of reverse transcriptase, thus creating the basis for one type of RT to outcompete the other ones within a given DGR system. Fig. 7 demonstrates proportions of six major RT clades in DGR systems in eighteen selected putative defense systems types. Noteworthy, for 12 out of 18 systems in more than 90% of occurrences were found to contain a single RT type (belong to the same DGR clade as defined by [6]).

**Fig. 7.**
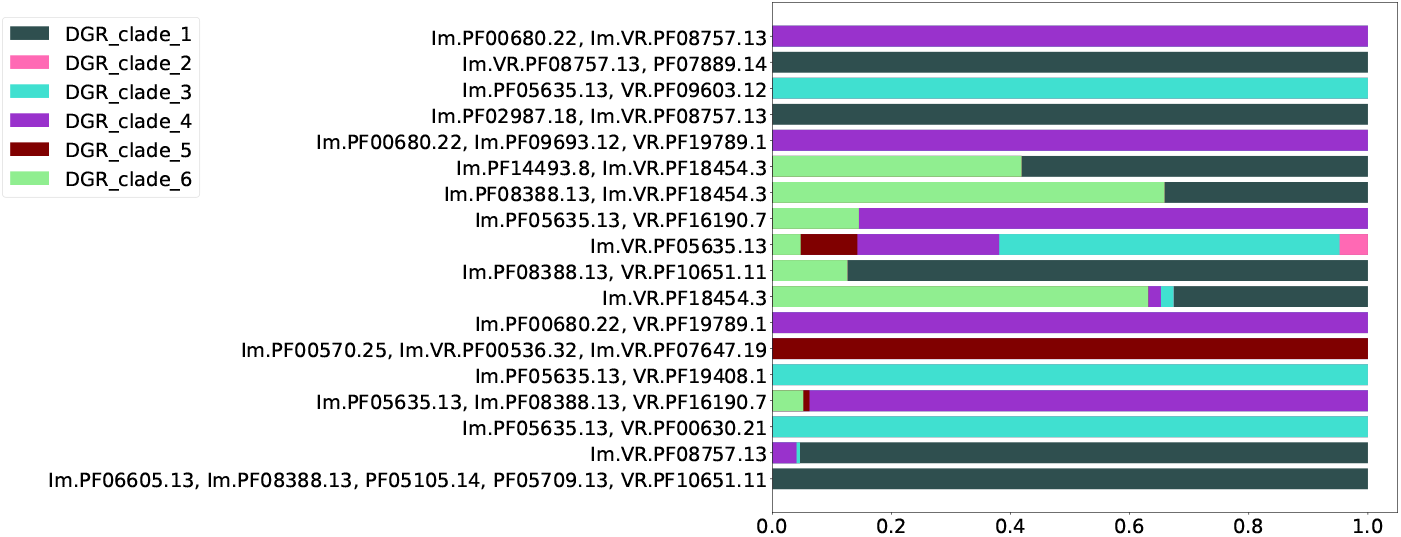
Proportion of major DGR clades based on RT phylogeny in eighteen selected Pfam compositions.

The biological significance of the classification of RT proteins of DGR systems into six major clades may correspond to the taxonomy of the organisms possessing DGR systems to a certain extent. One would expect that within the same taxon there should be DGR systems from the same RT clade. Alternatively, a DGR system of one type can be found in different non-related taxa, since a successful event of Horizontal Gene Transfer, once occurred, would establish a new lineage of DGR system within a recipient taxon. Thus, we expected to observe a distribution of Pfam compositions that is qualitatively similar to the distribution across major DGR clades. Fig. 8 shows distribution of the selected putative DGR-based defense systems among major prokaryotic taxa. Fig. 9 demonstrates that major prokaryotic taxa are strongly biased towards one or two major DGR clades corresponding to the selected Pfam compositions.

**Fig. 8.**
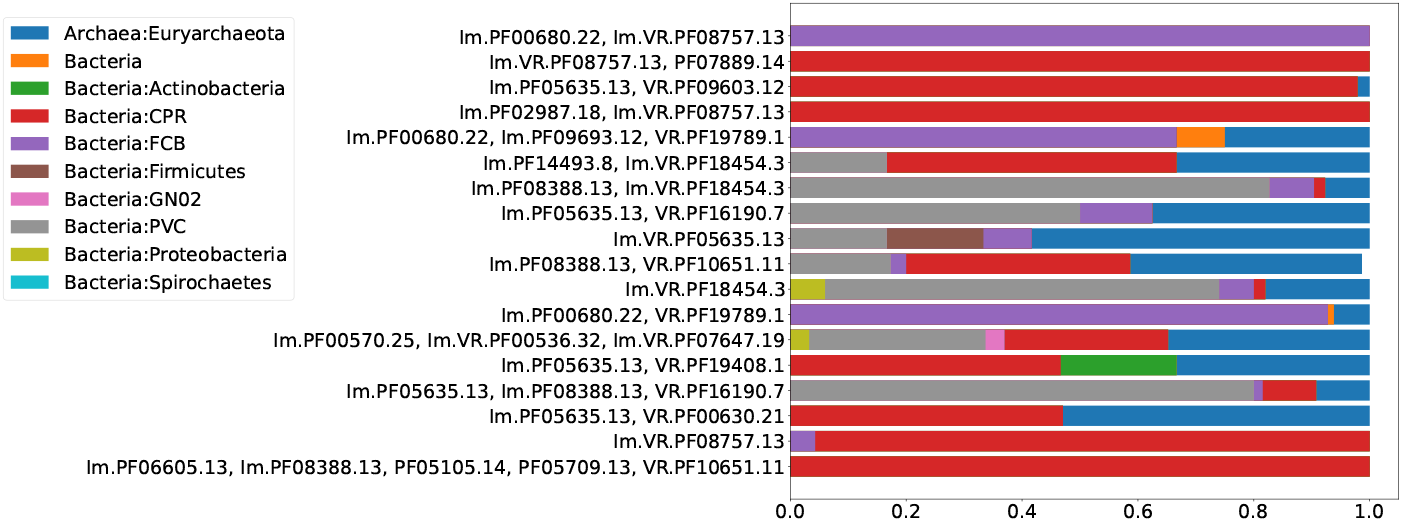
Distribution of eighteen selected Pfam compositions among major prokaryotic taxa.

**Fig. 9.**
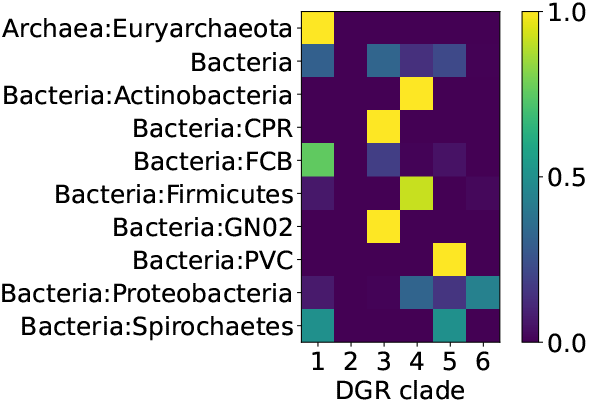
Proportions of major DGR clades within major prokaryotic taxa for eighteen selected Pfam compositions.

*·*Functional genetic systems typically demonstrate a more or less conserved order of genes, *e*.*g* see [20] and [22]. A set of *n* Pfams can be organized in *n*! possible permutations, and each Pfam can be encoded by one of two complementary strands giving 2*n* variants. Since the order of genes in an entire system can arbitrarily be chosen between forward and downstream order, the total number should be divided by a factor of two. Thus, *n* Pfams would yield 2^*n*^−1 *n*! possible arrangements, which translates to four if *n* equals two, and twenty four if *n* equals 3. One would expect observing a limited number of possible variants for order of genes, in the case of functional defense systems, with a fraction consisting the majority. Fig. 10 shows proportions of actual Pfam orders related to possible variant numbers for the eighteen selected compositions. Numbers in brackets show the number of given compositions.

**Fig. 10.**
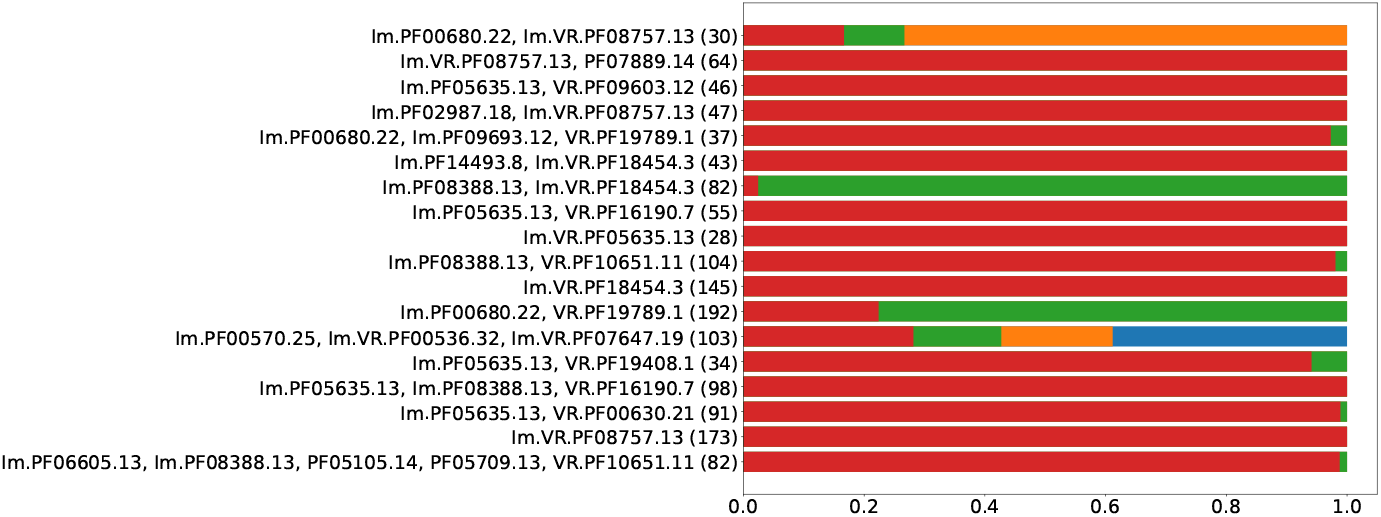
Proportions of gene arrangements among selected Pfam compositions. Numbers in brackets show amount of sequences containg given Pfam composition.

Finally we show representatives for four Pfam compositions containing more than two Pfams (Fig. 11). Majority of Pfams within these four compositions are poorly characterized or belong to domains with unknown function. In all four cases open reading frames are organized in compact operon-like modules.

**Fig. 11.**
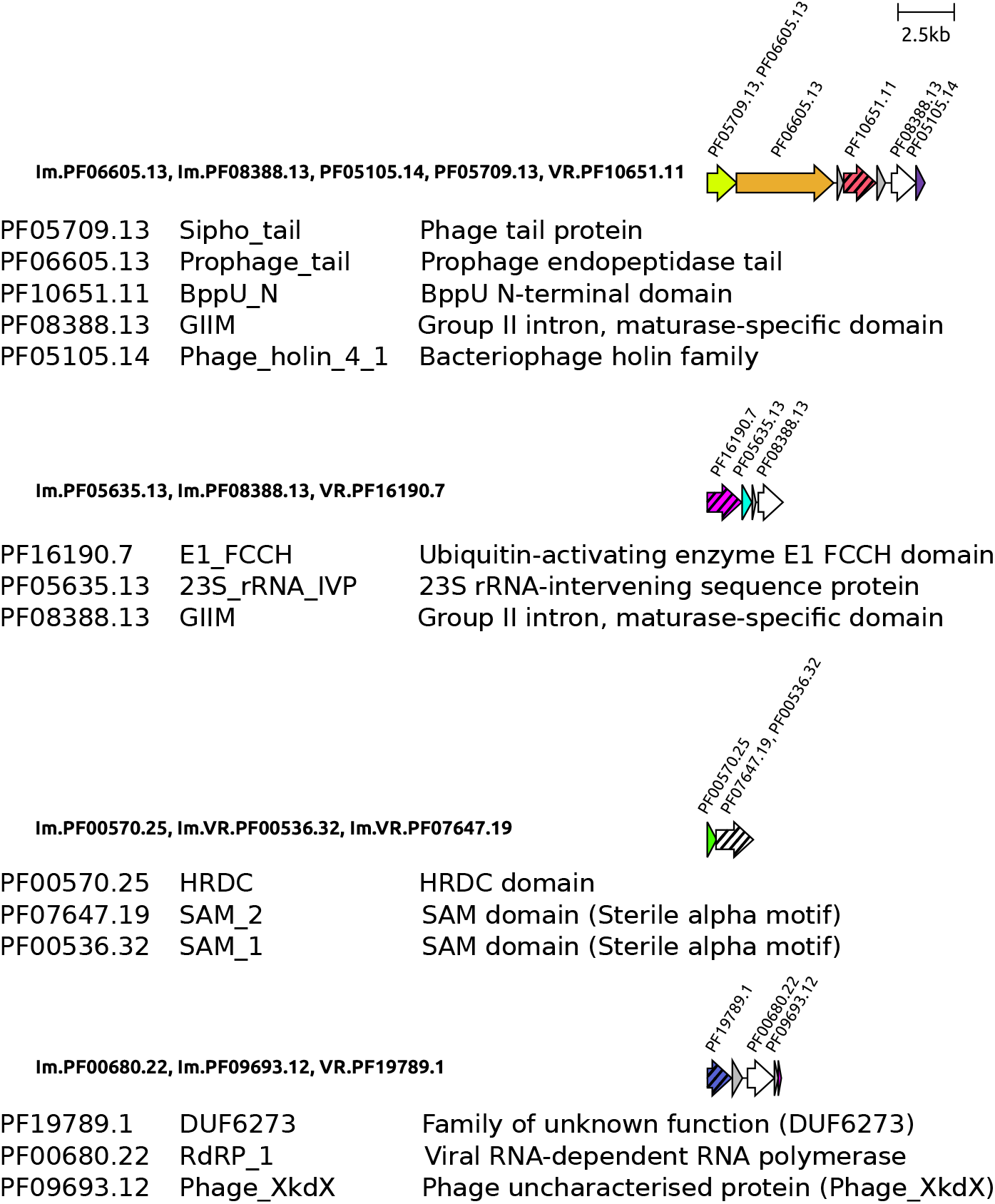
Organization of selected Pfam compositions. Open reading frames are shown as arrows. Reverse Transcriptase genes are shown in white. Hatching indicates proteins containing Variable Repeats.

## Methods

We have used all publicly available sequence data on DGR systems reported in [6, 14, 15]. We downloaded the nucleotide sequences encoding DGR systems along with the annotations from IGM and NCBI [23–25]. Ecological and taxonomic data were retrieved from [6].

To detect conserved protein domains (Pfams), that are potentially functionally linked to the components of DGR systems, we used modified Icity protocol [16]. This tool implements the ‘guilt by association’ principle, i.e. the preposition that functionally related genes often co-localise in the genomes (for example, genes, involved in a specific metabolic pathway, are frequently found in the same operon). co-localisation of genes in a sequence dataset may happen due to pure chance, depending on the amount of the data and possible overrepresentation of some organisms. These factors require statistical corrections, which are implemented in the Icity algorithm. The Icity protocol performs as the very first step the clustering of the protein sequences according to their similarity using a stochastic approach [26]. We replaced this step with clustering according to the Pfam families to which the protein domains belong [27]. The rationale behind the first step replacement is to further use data regarding Pfams associated with known prokaryotic defense machinery (see below). The proteins containing VR were analyzed separately from their non-VR peers from the same Pfam clusters.

All of the downstream analyses were performed according to the original Icity protocol. We applied the Icity protocol to each of the DGR components (RT, TR, and VR) separately. As a result we obtained protein clusters with associated Icity score, a metric reflecting the probability of functional association between given protein and the gene of interest (in this study the components of DGR systems). For further analysis we considered only the domains with the Icity scores greater than 0.7, following the recommendations of the authors of Icity protocol [16]. As a result we obtained a number of gene compositions, where components of DGR systems functionally interact with other genes with high confidence.

We next focused on gene compositions, where at least one protein family (domain) was known or predicted to be involved in prokaryotic antiviral defense, according to the recent large-scale studies [12, 13]. We visualized selected gene sets with upsetplot [28]. Proteins containing VR were treated as a separate entity from other sequences within the same Pfam cluster.

Codon usage bias was determined for all the open reading frames longer than 360 nucleotides within the putative DGR-systems ±10 kbp and for the region of the same size in the most distant part of the same sequence (assuming the sequence is circular). Sequences for which the distance between these two regions was less than 20 kbp were omitted from this analysis. Codon usage bias was estimated with the program chips from the EMBOSS package [29, 30].

Selected representatives of highly presented types of gene compositions, and containing more than two Pfams, were visualized with clinker [31]. We used R 4.1.2, python 3.8 and biopython [32] for data handling and visualization.

## Discussion

Roux and colleagues have demonstrated that DGR systems likely shaped virushost interactions in multiple taxa and biomes, representing a fundamental mechanism by which viruses and microbes adapt to a permanently changing environment [6]. Furthermore, DGR systems introduce more than 10% of all amino acid substitutions in some organisms. This level of diversity may preemptively adapt defense genes to yet-unknown potential invaders, as in the case of antibodies and T-cell receptors in vertebrates. In this study we demonstrated that several types of DGR systems co-localise and therefore are likely to functionally interact with known or predicted families of defense proteins [12, 13]. These gene systems pass all of the tests we applied for identifying functional modules. The results of our bioinformatic analysis enable identification of the most promising candidate systems for experimental evaluation of DGR-associated antiviral activity. Success rate in analogous attempts of new prokaryotic defense systems identification was 10 out of 28 tested candidates, and 29 out of 40 as reported in [12, 13], respectively.

Initiating this work we hypothesized a functional mechanism for DGR-driven immunity that would be analogous to mammalian adaptive immunity, wherein the recognition of the antigen particle by an immunoglobulin or T-cell receptor is followed by physical removal or inactivation of the target. Indeed, it is known that the VRs of some DGR systems are located in surface-exposed proteins [7] that are believed to serve as bacterial adhesins. However, the phage-specific sequestration of the phage virions by tight binding in a conformation unsuitable for infection may be one of the ways to reduce phage-induced mortality. The periplasmic localisation of the immune protein may provide an opportunity to interact with viral proteins involved in conduction of the DNA to interfere with the viral genome internalization. Interestingly, such mechanisms of the superinfection exclusion were described in phage T4 [33] and in phage HK97 [34] though without any connection to DGRs. Alternatively, binding of the viral proteins by variable cytosolic VR containing pathogen recognition proteins may be coupled with the directing of the target to degradation machinery as it is performed by ATP-dependent proteases adaptor proteins [35] or to direct the attack of the viral nucleo-proteid complexes, such as replication initiation complexes, that would lead to the degradation of the phage DNA. However, our data currently provide no hints for a possible mechanism of action of a DGR-based bacterial antiviral system. The experimental identification of the functional DGR-based immune systems among the predicted candidates would allow elucidating the underlying mechanism.

## Acknowledgments

We are grateful to Dr. Eugene E. Kulikov for helpful comments, Dr. Bob Blasdel Reuter and to Dr. Andrew Millard for critical reading and linguistic correction of the manuscript. We are also grateful to Alexandra Elbakyan for Sci-Hub.

## Declarations

### Funding

This work was supported by the Ministry of Education and Science of the Russian Federation.

### Conflict of interest/Competing interests (check journal-specific guidelines for which heading to use)

The authors declare no competing interests.

### Ethics approval

NA

### Consent to participate

NA

### Consent for publication

NA

### Availability of data and materials

All the data we used are publicly available.

### Code availability

Python code reproducing our results will be publicly available immediately upon the manuscript acceptance for publication.

### Authors’ contributions

I.B. formulated the hypothesis and designed the study. A.S. and I.B. performed the analyses. All authors contributed to and commented on results interpretation. I.B. prepared the manuscript and figures. A.L. devised original idea. All authors reviewed, contributed to improving, and approved the initial manuscript and figures.

## Notes

### Competing Interest Statement

The authors have declared no competing interest.

